# The effects of *trans*-chalcone and chalcone hydrate on the growth of *Babesia* and*Theileria*

**DOI:** 10.1101/480657

**Authors:** Gaber El-Saber Batiha, Amany Magdy Beshbishy, Dickson Stuart Tayebwa, Oluyomi Stephen Adeyemi, Hazem Shaheen, Naoaki Yokoyama, Ikuo Igarashi

## Abstract

**Background:** Chemotherapy is a principle tool for the control and prevention of piroplasmosis. The search for a new chemotherapy against *Babesia* and *Theileria* parasites has become increasingly urgent due to the toxic side effects of and developed resistance to the current drugs. Chalcones have attracted much attention due to their diverse biological activities. With the aim to discover new drugs and drug targets, *in vitro* and *in vivo* antibabesial activity of *trans*-chalcone (TC) and chalcone hydrate (CH) alone and combined with diminazene aceturate (DA), clofazimine (CF) and atovaquone (AQ) were investigated.

**Methodology/Principal findings:** The fluorescence-based assay was used for evaluating the inhibitory effect of TC and CH on five of *Babesia* and *Theileria* species, including *B. bovis*, *B. bigemina*, *B. divergens*, *B. caballi*, and *T. equi*, the combination with DA, CF, and AQ on *in vitro* cultures, and on the multiplication of a *B. microti*–infected mouse model. The cytotoxicity of compounds was tested on Madin– Darby bovine kidney (MDBK), mouse embryonic fibroblast (NIH/3T3), and human foreskin fibroblast (HFF) cell lines. The half maximal inhibitory concentration (IC_50_) values of TC and CH against *B. bovis*, *B. bigemina*, *B. divergens*, *B. caballi*, and *T. equi* were 69.6 ± 2.3, 33.3 ± 1.2, 64.8 ± 2.5, 18.9 ± 1.7, and 14.3 ± 1.6 µM and 138.4 ± 4.4, 60.9 ± 1.1, 82.3 ± 2.3, 27.9 ± 1.2, and 19.2 ± 1.5 µM, respectively. In toxicity assays, TC and CH affected the viability of MDBK, NIH/3T3, and HFF cell lines the with half maximum effective concentration (EC_50_) values of 293.9 ± 2.9, 434.4 ± 2.7, and 498 ± 3.1 µM and 252.7 ± 1.7, 406.3 ± 9.7, and 466 ± 5.7 µM, respectively. In the mouse experiment, TC reduced the peak parasitemia of *B*. *microti* by 71.8% when administered intraperitoneally at 25 mg/kg. Combination therapies of TC–diminazene aceturate and TC–clofazimine were more potent against *B. microti* infection in mice than their monotherapies.

**Conclusions/Significance:** In conclusion, both TC and CH inhibited the growth of *Babesia* and *Theileria in vitro*, and TC inhibited the growth of *B. microti in vivo.* Therefore, TC and CH could be candidates for the treatment of piroplasmosis after further studies.

**Author summary:** Protozoa of the genus *Babesia* are the second most common blood-borne parasites of mammals after the trypanosomes. *Babesia* and *Theileria* are the etiological agents of piroplasmosis, a tick-transmitted disease causing substantial losses of livestock and companion animals worldwide and has recently gained attention as one of the emerging zoonosis in humans. Diminazene aceturate and imidocarb dipropionate are still the first choices for the treatment of animals. However, these drugs cause many adverse effects. Furthermore, they are not approved for human medicine. Therefore, the development of alternative treatment remedies against babesiosis is urgently required. In the present study we evaluated the effects chalcone hydrate (CH) and *trans*-chalcone (TC), against the growth of four species of *Babesia* and *T. equi.* Furthermore, we studied the chemotherapeutic potential of TC on *B. microti* in mice. The effects of the combined treatment of TC with DA, CF and AQ revealed that TC was found to diminish the adverse effects of these drugs

## 1. Introduction

Babesiosis is one of the most severe infections of animals worldwide and has recently gained attention as one of the emerging zoonosis in humans [1, 2]. *Babesia bovis*, *Babesia bigemina*, and *Babesia divergens* infect cattle and cause bovine babesiosis. Of these, *B. bovis* is much more virulent than *B. bigemina* and *B. divergens* due to its ability to sequestrate in the capillaries, causing hypotensive shock syndrome and neurological damage [3]. In horses, *Babesia caballi* and *Theileria equi* (formerly *Babesia equi*) infect horses, causing equine piroplasmosis. *T. equi* parasitizes leucocytes and erythrocytes for the completion of its life cycle, causing anemia, weight loss, lethargy, and fever [4], whereas *B. caballi* directly infects horse erythrocytes in a manner similar to *B. bovis* and *B. bigemina* in cattle. Human babesiosis is caused by *Babesia microti* in North America, while in Europe, it is caused by *Babesia divergens*. Human babesiosis manifests as an apparently silent infection to a fulminant, malaria-like disease, resulting occasionally in the death of the infected individual [5].

Prevention of babesiosis relies on vector control, vaccination, and chemotherapy. Thus far, chemotherapy has been the most successful method due to the availability of efficacious compounds such as diminazene aceturate and imidocarb dipropionate for animals and atovaquone, azithromycin clindamycin, and quinine for humans [5]. Unfortunately, atovaquone-resistant *Babesia gibsoni* has been reported [6, 7], and Mosqueda et al. (2012) reported the emergence of parasites resistant to diminazene aceturate (DA) [8]. Therefore, research to discover new drugs and drug targets is the fundamental approach toward addressing current limitations.

Chalcones (*trans*-1, 3-diaryl-2-propen-1-ones) are natural products belonging to the flavonoid family that are widespread in plants and are considered as intermediate in the flavonoid biosynthesis [9, 10]. They are recognized for their broad-spectrum biological activities, including antimalarial [11], anticancer, antileishmanial, antitrypanosomal [12, 13, 14, 15, 16], anti-inflammatory, antibacterial, antioxidant, antifilarial, antifungal, antimicrobial, larvicidal, and anticonvulsant ones [17, 18]. Based on the wide range of pharmacological effects, it is implied that chalcones have several modes of action in different parasites. For instance, Go et al. (2004) showed that chalcones modulate the permeability pathways of the *Plasmodium*-infected erythrocyte membrane, affecting its growth and multiplication [9]. Frölich et al. (2005) showed that chalcones inhibit glutathione (GSH)-dependent haemin degradation and binding to the active site of the cysteine protease (falcipain) enzyme involved in hemoglobin degradation in the *Plasmodium* parasite [11, 19, 20]. Chalcones inhibit the components of mitochondrial respiratory chain *bc_1_* complex (ubiquinol–cytochrome c reductase) (UQCR) [21]. Additionally, chalcones inhibit the cyclin-dependent protein kinases (CDKs) (Pfmrk and PfPK) and plasmepsin II [22]. These various pathways demonstrate the importance of chalcones as chemotherapeutic candidates against malaria. However, the effect of chalcones had never been evaluated against *Babesia* and *Theileria* parasites. Therefore, this study evaluated the effects of chalcones, namely, chalcone hydrate (CH) and *trans*-chalcone (TC), against the growth of *B. bovis*, *B. bigemina*, *B. divergens*, *B. caballi*, and *T. equi in vitro*. Furthermore, we studied the chemotherapeutic potential of TC on *B. microti* in mice.

## 2. Materials and methods

### 2.1. Cultivation conditions

#### 2.1.1. Parasites and mice

The German bovine strain of *B. divergens* [23], the Texas strain of *B. bovis*, the Argentina strain of *B. bigemina*, and the United States Department of Agriculture (USDA) strains of *T. equi* and *caballi* were used for the *in vitro* studies, while the Munich strain of *B. microti* was used for the *in vivo* studies [24]. To perform the *in vivo* studies, female BALB/c mice (CLEA Japan, Inc., Tokyo) housed under a pathogen-free environment with controlled temperature (22^º^C) and humidity and a 12 h light/dark cycle were used for the cultivation of *B. microti*.

#### 2.1.2. Reagents and chemicals

The *trans*-chalcone (TC) **(Fig. 1A)**, chalcone hydrate (CH) **(Fig. 1B)**, diminazene aceturate (DA), clofazimine (CF), and atovaquone (AQ) powders (Sigma-Aldrich Japan, Tokyo, Japan) were prepared in dimethyl sulfoxide (DMSO) (Wako Pure Chemical Industries, Ltd., Osaka, Japan) in 10 mM stock solutions and stored at −30ºC.

**Fig 1.**
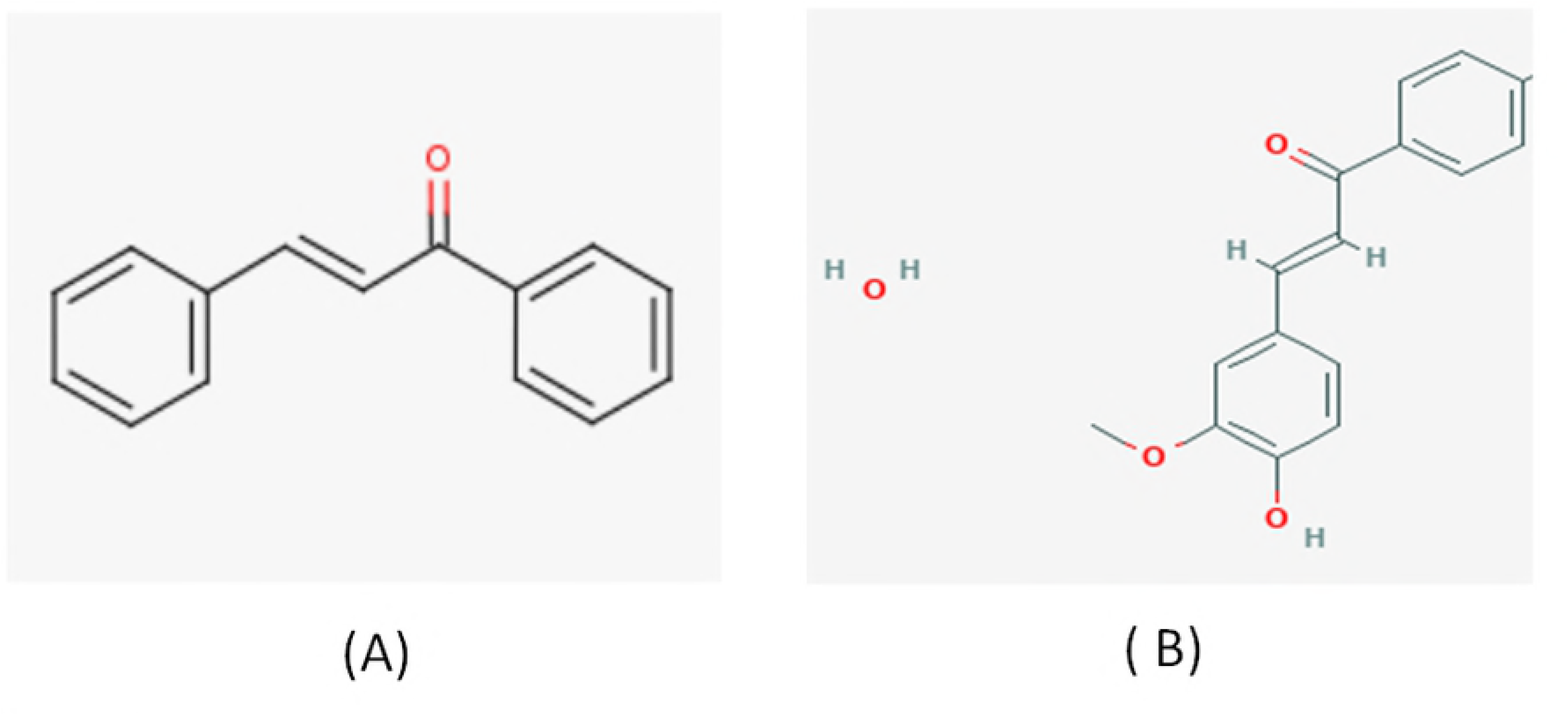
Chemical structure of *trans*-chalcone (A) and chalcone hydrate (B) used in this study

The 10,000×SYBR Green 1 (SG1) nucleic acid stain (Lonza America, Alpharetta, GA, USA) was purchased and stored at −30ºC, wrapped in aluminum foil paper for protection from direct light. A lysis buffer containing Tris (130 mM at pH 7.5), EDTA (10 mM), saponin (0.016% w/v), and Triton X – 100 (1.6% v/v) was prepared and stored at 4ºC.

#### 2.1.3. Cultivation of parasites *in vitro*

The purified bovine red blood cells (RBCs) were used to maintain *B. bovis*, *B. bigemina*, and *B. divergens*, and the purified equine RBCs were used to maintain *B. caballi* and *T. equi*. The cultivation was performed in the micro-aerophilic stationary-phase culture system at 37ºC, 5% CO_2_, 5% O_2_, and 90% N_2_ as previously described [23, 24]. M199 (Sigma-Aldrich, Tokyo, Japan), supplemented with 40% bovine serum was used to culture *B. bigemina* and *B. bovis*. Medium RPMI 1460 (Sigma-Aldrich, Tokyo, Japan) supplemented with 40% bovine serum was used to culture *B. divergens*. M199 supplemented with 40% equine serum and 13.6 µg/mL of hypoxanthine (MP Biomedicals, Santa Ana, CA, USA) was used to cultivate *T. equi*. GIT medium (Sigma-Aldrich, Tokyo, Japan) supplemented with 40% equine serum was used to maintain the *B. caballi* culture. To prevent bacterial and fungal contamination, 60 µg/mL of streptomycin and 0.15 µg/mL of amphotericin B (Sigma-Aldrich Corp., St. Louis, MO, USA) were added to all of the culture media.

### 2.2. *In vitro* growth inhibitory effects

The half maximal inhibitory concentration (IC_50_) for CH, TC, DA, AQ, and CF was determined using the fluorescence assay as previously described [23]. Briefly, 12.5, 25, 50, 100, and 200 µM CH and TC and 0.01, 0.1, 1, 10, and 100 μM DA, AQ, and CF were used to determine the inhibition concentration in a 96-well plate with a 2.5% hematocrit for *B. bovis* and *B. bigemina* and a 5% hematocrit for *B. divergens*, *B. caballi*, and *T. equi*. The plates were cultivated for 4 days without changing the media. On day 4, 100 µL of lysis buffer containing 2×SG1 was directly added to each well and gently mixed by pipetting. The plate was wrapped in aluminum foil paper for protection from direct light and incubated for 6 h at room temperature. After that, the plates were placed into the fluorescence spectrophotometer (Fluoroskan Ascent; Thermo Scientific, San Diego, CA, USA). The relative fluorescence values were read at 485 and 518 nm excitation and emission wavelengths, respectively. Positive control wells containing infected red blood cells (iRBCs) and uninfected red blood cells (RBCs) were included as negative control wells. Gain values were set to percentages after subtraction of the mean values of the negative control and transferred into GraphPad Prism (GraphPad Software, Inc., San Diego, California, USA) to calculate the IC_50_ value using the non-linear regression analysis (curve fit).

### 2.3. Morphological changes and viability experiment in vitro

The microscopy assay was performed as previously described [24]. Five concentrations at 0.25×, 0.5×, 1×, 2×, and 4× the IC_50_ of CH, TC, and DA were used for this experiment. A 100 µL reaction volume containing 90 µL of respective media and 10 µL of iRBCs normalized to 1% parasitemia was incubated in a 96-well microtiter plate at 37ºC in a humidified multi-gas water-jacketed incubator. The 90 µL of media was changed daily and replaced with 90 µL of new media containing the same concentration of drugs (CH or TC) for 4 consecutive days. In the course of the 4 days of treatment, Giemsa-stained thin blood smears were prepared, and the parasitemia in 10,000 RBCs was monitored every 24 h. On the 5^th^ day, 3 µL of RBCs from each well was mixed with 7 µL of fresh RBCs, transferred into a new 96-well microtiter plate, and cultured in drug-free media. The media were replaced every day, and the parasitemia was monitored every 2 days until 6 days after the last treatment. The viability of drug-treated parasites was checked in the blood smear for 6 days after the last treatment. The presence of parasites was recorded as positive (relapse), and the absence of parasites was recorded as negative (total parasite clearance). Each experiment was performed in triplicate in three separate trials. The morphological changes were observed under a light microscope, and micrographs were captured using Nikon Digital Sight ^®^ (Nikon Corporation, Japan).

### 2.4. Combination treatment using CH or TC with DA, AQ, or CF in vitro

The combination studies were performed in accordance with the previously described protocol [24]. Three sets of duplicate wells with five selected concentrations of CH, TC, DA, AQ, and CF at 0.25×, 0.5×, 1×, 2×, and 4× the IC_50_ were loaded in a 96-well plate. The first set of wells contained concentrations of single CH treatments, the second set contained concentrations of single DA or AQ or CF treatment, and the third set contained the combination of CH with DA, AQ, or CF at a constant ratio (1:1). The same experiment was repeated for TC with DA, AQ, and CF in three separate trials. The cultivation was performed for 4 days in a 100 µL reaction volume of media containing the drug concentrations and a hematocrit of 2.5% for *B. bovis* and *B. bigemina* and 5% for *B. divergens*, *B. caballi*, and *T. equi*. On day 4, 100 µL of lysis buffer containing 2×SG1 was added. The plate was wrapped with aluminum foil for protection from light and incubated at room temperature for 6 h. The plates were then loaded into a fluorescence spectrophotometer, and the relative fluorescence values were read at 485 and 518 nm excitation and emission wavelengths, respectively. The obtained fluorescence values were set to percentages after subtraction of the mean values of the negative control. The growth inhibition values obtained were entered into CompuSyn software^®^ (ComboSyn, Inc., Paramus, NJ, USA) [25] for calculation of the degree of association based on the combination index (CI) values. The CI values of the drug combination were determined using the formula [(1×IC_50_) + (2×IC_75_) + (3×IC_90_) + (4×IC_95_)]/10, and the drug combination was considered synergistic if the value was less than 0.90, additive if the value was 0.90–1.10, and antagonistic if the value was more than 1.10 [25].

### 2.5. Evaluation of the effects of CH and TC on the host erythrocyte in vitro

The effects of CH and TC on the host RBCs (bovine and equine) were investigated as previously described [24]. Bovine and equine RBCs were incubated in the presence of 10, 100, and 200 µM CH and TC for 3 and 6 h at 37ºC. The RBCs were then washed three times with drug-free media and subsequently used for the cultivation of *B. bovis* and *T. equi.* The effect was monitored using the fluorescence assay.

### 2.6. Cell cultures

Madin–Darby bovine kidney (MDBK), mouse embryonic fibroblast (NIH/3T3), and human foreskin fibroblast (HFF) cell lines were cultured continuously at 37ºC in a humidified incubator with 5% CO_2_. MDBK cell line was maintained in 75 cm^2^ culture flasks with Minimum Essential Medium Eagle (MEM, Gibco, Thermo Fisher Scientific, Carlsbad, CA, USA), while Dulbecco’s Modified Eagle’s Medium (DMEM, Gibco, Thermo Fisher Scientific, Carlsbad, CA,USA) was used for NIH/3T3, and HFF cell lines cultivation. Each medium was supplemented with 10% fetal bovine serum, 0.5% penicillin/streptomycin (Gibco, Thermo Fisher Scientific, Carlsbad, CA, USA), and an additional 1% glutamine. The medium was changed every 2 to 4 days and incubated until approximately 80% confluent. The cells were checked by staining with 4, 6-diamidino-2-phenylindole dihydrochloride (DAPI; Sigma-Aldrich Corp., St. Louis, MO, USA) to ensure free mycoplasma contamination. TrypLETM Express (Gibco, Thermo Fisher Scientific, Carlsbad, CA, USA) was used to allow cell detachment from the culture flask after washing two times with Dulbecco’s phosphate-buffered saline (DPBS). Subsequently, viable cells were stained with 0.4% trypan blue solution and then counted using a Neubauer improved C-Chip (NanoEnTek Inc., Seoul, Korea).

### 2.7. Cytotoxicity assay of CH, TC, DA, AQ, and CF on MDBK, NIH/3T3, and HFF cell lines

The drug-exposure viability assay was performed in accordance with the recommendation for the cell-counting Kit-8 (CCK-8, Dojindo, Japan). In a 96-well plate, 100 µL of cells at a density of 5×10^4^ cells/mL was seeded per well and allowed to attach to the plate for 24 h at 37℃ in a humidified incubator with 5% CO_2_. For CH and TC, 10 µL of twofold dilutions was added to each well to a final concentration of 12.5–500 µM in triplicate, while for DA, AQ, and CF, 10 µL of twofold dilutions was added to each well to a final concentration of 100 µM in triplicate. The wells with only a culture medium were used as blanks, while the wells containing cells and a medium with 0.4% DMSO were used as a positive control. The exposure of drugs was carried out for 24 h, followed by the addition of 10 µL of CCK-8. The plate was further incubated for 3 h, and the absorbance was measured at 450 nm using a microplate reader.

### 2.8. The chemotherapeutic effect of TC against *B. microti* in mice

The *in vivo* inhibitory effects of TC were evaluated against *B. microti* in mice as previously described [23]. Briefly, 50 female BALB/c mice at 8 weeks of age were caged in 10 groups (five mice/group). *B. microti* recovered from the frozen stock (stored at −80ºC) was thawed and injected into two mice intraperitoneally. The parasitemia was monitored daily via microscopy. The mice were sacrificed and blood collected by cardiac puncture when the parasitemia was over 40%. The blood was diluted with phosphate-buffered saline to obtain an inoculum containing 1×10^7^/mL of *B. microti* iRBCs. The mice in groups 2–10 received 0.5 mL intraperitoneal (IP) injections of the inoculum (1×10^7^ *B. microti* iRBCs). Group 1 was left uninfected and untreated as a control. When the average parasitemia in all mice reached 1%, drug treatment was initiated for 5 days. The mice in group 2 received sesame oil via IP injection as a control. Groups 3–6 received a 25 mg/kg IP injection of TC, a 25 mg/kg IP injection of DA, oral administration of 20 mg/kg of AQ, and oral administration of 20 mg/kg of CF, respectively. Groups 7–9 were treated with a combination of (12.5 mg/kg of TC and 12.5 mg/kg of DA), (12.5 mg/kg of TC and 10 mg/kg of AQ), and (12.5 mg/kg of TC and 10 mg/kg of CF), respectively, via a route similar to that for the single drug, while the mice in group 10 received DDW via IP injection as a control (infected and untreated mice). The parasitemia and blood parameters were monitored via microscopy and a hematology analyzer (Celltac α MEK-6450, Nihon Kohden Corporation, Tokyo, Japan) every 2 and 4 days, respectively. The experiment was repeated twice. On day 45, blood was collected for PCR detection of the parasites.

### 2.9. The genomic DNA extraction and PCR detection of *B. microti*

Genomic DNA was extracted from the blood using a QIAamp DNA Blood Mini Kit (Qiagen, Tokyo, Japan). A nested PCR (nPCR) targeting a small-subunit rRNA (ss-rRNA) gene in *B. microti* was performed as described previously [24]. Briefly, the PCR amplifications were performed in a 10 µL reaction mixture containing 0.5 µM of each primer, 0.2 mM dNTP mix, 2 µL of 5× SuperFi™ buffer, 0.1 µL of Platinum SuperFi™ DNA polymerase (Thermo Fisher Scientific, Japan), 1 µL of DNA template, and 4.9 µL of DDW. The cycling conditions were 94ºC for 30 s denaturation, 53ºC for 30 s annealing, and 72ºC for 30 s as extension steps for 35 cycles, using the forward (5′-CTTAGTATAAGCTTTTATACAGC-3′) and reverse (5′-ATAGGTCAGAAACTTGAATGATACA-3′) primers. Afterward, 1 µL of the DNA template from the first PCR amplification was used as the template for the nPCR assays under similar cycling conditions, using the forward (5′-GTTATAGTTTATTTGATGTTCGTTT-3′) and reverse (5′-AAGCCATGCGATTCGCTAAT-3′) primers. After that, the PCR products were determined by electrophoresis in a 1.5% agarose gel, stained with ethidium bromide, and visualized under the UV Tran illuminator. The bands with an expected size of 154 bp were considered positive.

### 2.10. Statistical analysis

The IC_50_ values of CH, TC, DA, AQ, and CF were determined using the non-linear regression curve fit in GraphPad Prism (GraphPad Software, Inc., USA). The difference in parasitemia, hematology profile, and body weight was analyzed using an independent student’s *t*-test. A *p*-value < 0.05 was considered statistically significant.

### 2.11. Ethical clearance

All experiments were approved by the Animal Care and Use Committee and conducted in accordance with Regulations on Management and Operation of Animal Experiments as stipulated by Obihiro University of Agriculture and Veterinary Medicine (accession number of animal experiment: 28-111-2/28-110). These regulations were established by Fundamental Guidelines for Proper Conduct of Animal Experiment and Related Activities in Academic Research Institutions, the Ministry of Education, Culture, Sports and Technology (MEXT), Japan.

## 3. Results

### 3.1. The growth inhibitory effect of chalcones against *Babesia* and *Theileria*

The growth inhibitory assay was conducted on five species: *B. bovis*, *B. bigemina*, *B. divergens*, *B. caballi*, and *T. equi*. *Trans*-chalcone (TC) and chalcone hydrate (CH) inhibited the multiplication and growth of all species tested in a dose-dependent manner **(Figs 2 and 3)**.

**Fig 2.**
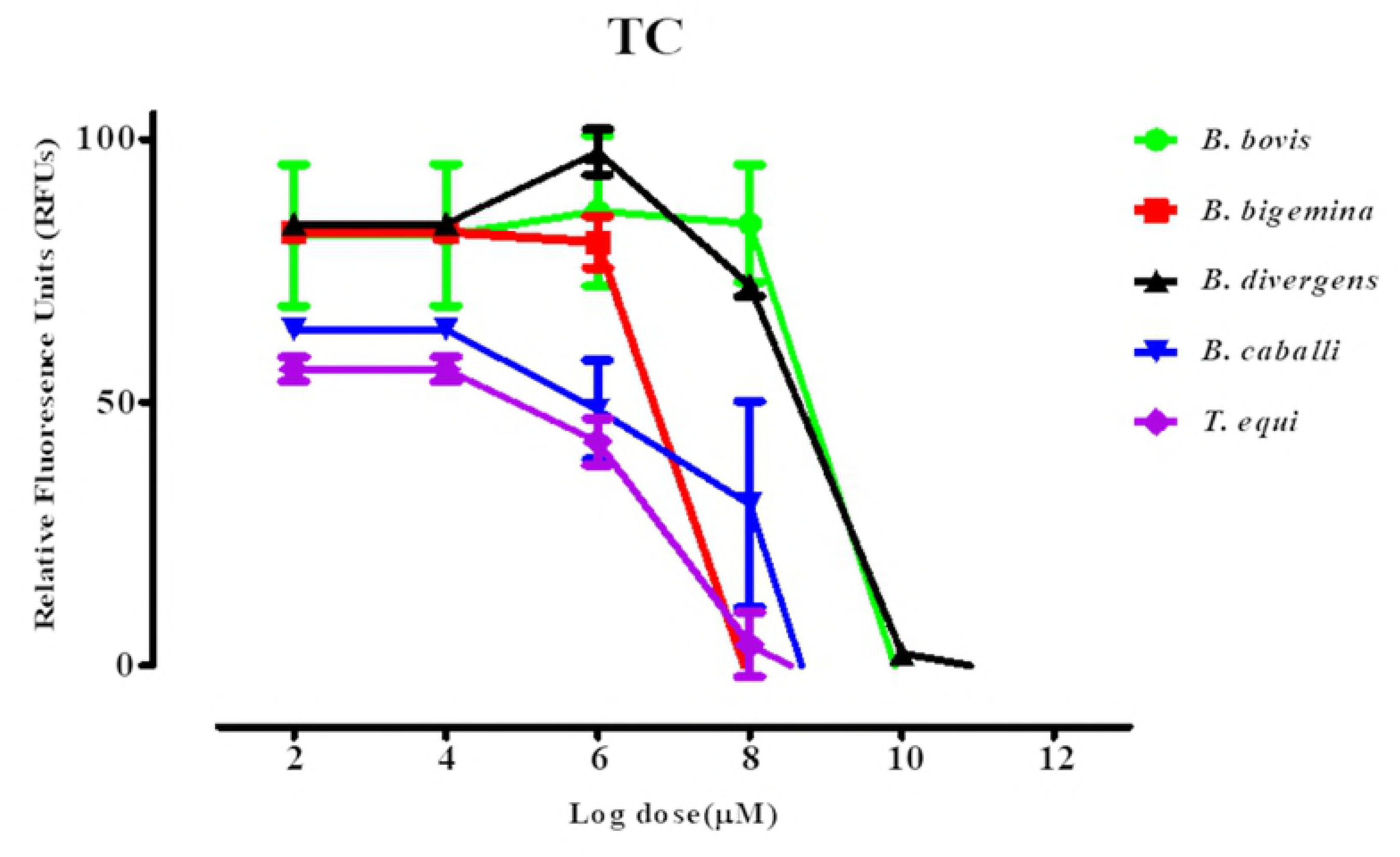
Dose-response curves of TC against *Babesia* and *Theileria* parasites *in vitro*. The curves show the relative fluorescence units of *B. bovis*, *B. bigemina*, *B. divergens*, *B. caballi*, and *T. equi* treated with increasing concentrations of TC. The results were determined via fluorescence assay after 96 h of incubation in three separate trials. The values obtained from three separate trials were used to determine the IC_50_ values using the non-linear regression (curve fit analysis) in GraphPad Prism software (GraphPad Software Inc., USA).

**Fig 3.**
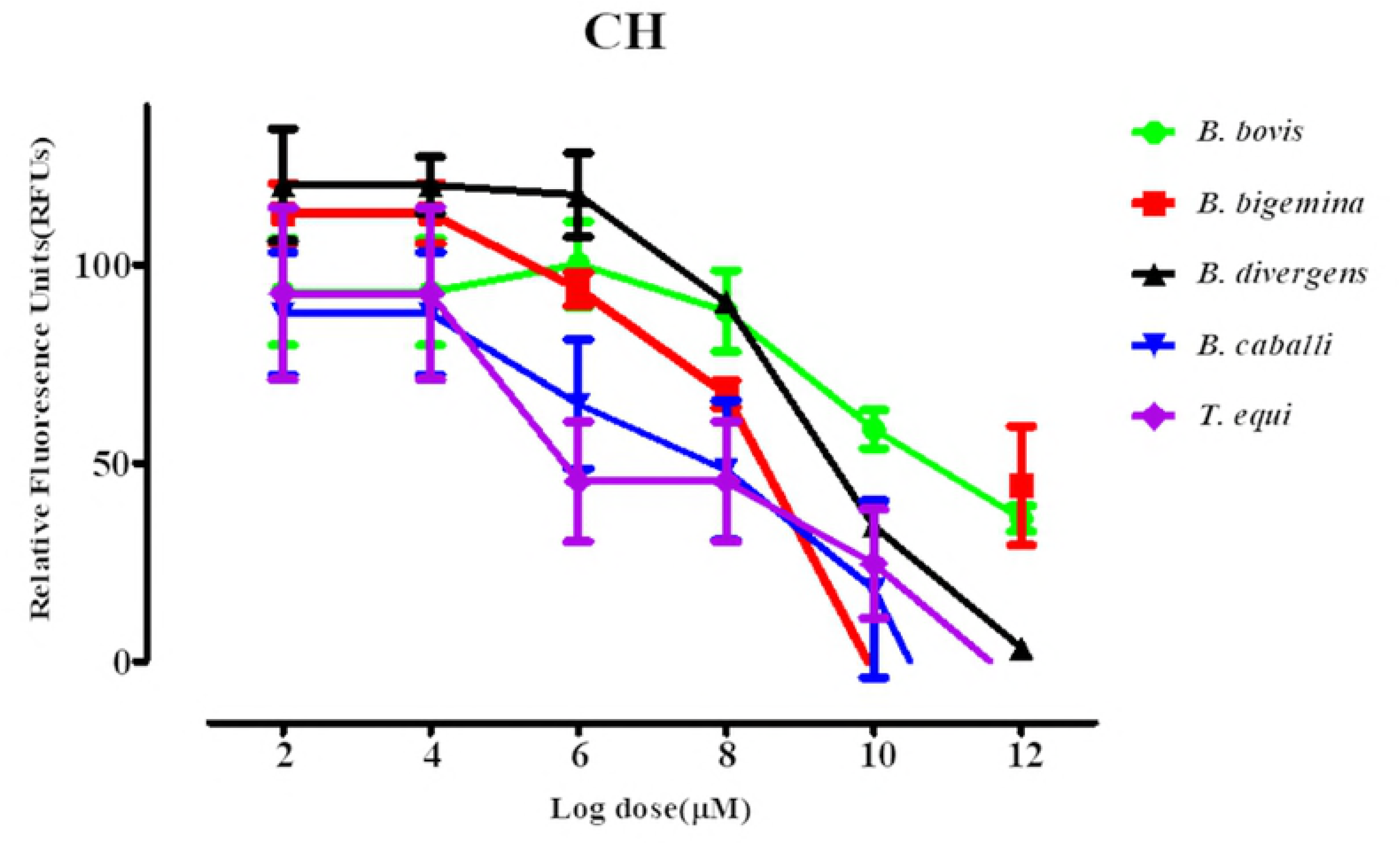
Dose-response curves of CH against *Babesia* and *Theileria* parasites *in vitro*. The curves show the relative fluorescence units of *B. bovis*, *B. bigemina*, *B. divergens*, *B. caballi*, and *T. equi* treated with increasing concentrations of CH. The results were determined via fluorescence assay after 96 h of incubation in three separate trials. The values obtained from three separate trials were used to determine the IC_50_ values using the non-linear regression (curve fit analysis) in GraphPad Prism software (GraphPad Software Inc., USA).

The IC_50_ values of TC and CH on *B. bovis*, *B. bigemina*, *B. divergens*, *B. caballi*, and *T. equi* were 69.6, 33.3, 64.8, 18.9, and 14.3 μM and 138.4, 60.9, 82.3, 27.9, and 19.2 μM, respectively (Table 1)

**Table 1.** The IC_50_ and selectivity index of TC and CH

| Compounds | Parasites | IC<sub>50</sub> (μM)<sup>a</sup><br>parasites | EC<sub>50</sub> (μM)<sup>b</sup> |  |  | Selective index<sup>c</sup> |  |  |
| --- | --- | --- | --- | --- | --- | --- | --- | --- |
|  |  |  | MDBK | NIH/3T3 | HFF | MDBK | NIH/3T3 | HFF |
| TC | *B. bovis* | 69.6 ± 2.3 | 293.9 ± 2.9 | 434.4 ± 2.7 | 498 ± 3.1 | 4.2 | 6.2 | 7.2 |
|  | *B. bigemina* | 33.3 ± 1.2 |  |  |  | 8.8 | 13.1 | 15.0 |
|  | *B. divergens* | 64.8 ± 2.5 |  |  |  | 4.5 | 6.7 | 7.7 |
|  | *B. caballi* | 18.9 ± 1.7 |  |  |  | 15.6 | 23.0 | 26.3 |
|  | *T. equi* | 14.3 ± 1.6 |  |  |  | 20.6 | 30.4 | 34.8 |
|  | *P. falciparum* | 11.5* |  |  |  |  |  |  |
| CH | *B. bovis* | 138.4 ± 4.4 | 252.7 ± 1.7 | 406.3 ± 9.7 | 466.3 ± 5.7 | 1.8 | 2.9 | 3.4 |
|  | *B. bigemina* | 60.9 ± 1.1 |  |  |  | 4.1 | 6.7 | 7.7 |
|  | *B. divergens* | 82.3 ± 2.3 |  |  |  | 3.1 | 4.9 | 5.7 |
|  | *B. caballi* | 27.9 ± 1.2 |  |  |  | 9.1 | 14.6 | 16.7 |
|  | *T. equi* | 19.2 ± 1.5 |  |  |  | 13.2 | 21.2 | 24.3 |
|  | *P. falciparum* | 21.7** |  |  |  |  |  |  |

<sup>a</sup> Half maximum inhibition concentration of *trans*-chalcone (TC) and chalcone hydrate (CH) on the *in vitro* culture of parasites. The value was determined from the dose-response curve using non-linear regression (curve fit analysis). The values are the means of triplicate experiments.
<sup>b</sup> Half maximum effective concentration of TC and CH on the cell line. The values were determined from the dose-response curve using non-linear regression (curve fit analysis). The values are the means of triplicate experiments.
<sup>c</sup> Ratio of the EC<sub>50</sub> of cell lines to the IC<sub>50</sub> of each species. High numbers are favorable.
\*(Geyer et al., 2009)
\*\* (Go et al., 2004)

In this study, DA showed IC_50_ values at 0.35, 0.68, 0.43, 0.022, and 0.71 µM against *B. bovis*, *B. bigemina*, *B. divergens*, *B. caballi*, and *T. equi*, respectively. AQ showed IC_50_ values at 0.039, 0.701, 0.038, 0.102, and 0.095 µM against *B. bovis*, *B. bigemina*, *B. divergens*, *B. caballi*, and *T. equi*, respectively. CF showed IC_50_ values at 8.24, 5.73, 13.85, 7.95, and 2.88 µM against *B. bovis*, *B. bigemina*, *B. divergens*, *B. caballi*, and *T. equi*, respectively **(S1 Table)**. The effectiveness of chalcones was not influenced by the diluent, since there was no significant difference in the inhibition between wells containing the DMSO and untreated wells. The precultivation of RBCs with TC and CH was conducted to determine their direct effect on host RBCs. Bovine and equine RBCs were incubated with TC or CH at 10, 100, and 200 µM for 3 h prior to the subculture of *B. bovis* and *T. equi*. The multiplication of *B. bovis* and *T.equi* did not signicantly differ between TC- or CH-treated RBCs and normal RBCs for either species (data not shown).

### 3.2. The viability of parasites treated with *trans*-chalcone and chalcone hydrate and the morphological changes in treated parasites

A viability assay was performed to determine whether the concentrations of TC and CH could completely clear parasites after 4 days of successive treatment, followed by withdrawal of the drug pressure. *B. bovis*, *B. bigemina*, and *B. caballi* treated with TC could not regrow at a concentration of 2×IC_50_, while *B. divergens* could not regrow at 4×IC_50_. *B. bovis*, *B. bigemina*, *B. divergens*, and *B. caballi* treated with CH could not regrow at a concentration of 4×IC_50_. *T. equi* treated with TC and CH could regrow at a concentration of 4×IC_50_ **(Table 2)**.

**Table 2.** The viability of *Babesia* and *Theileria* parasites treated with TC and CH. The positive symbol (+) indicates regrowth of the parasites, and the negative symbol (-) indicates total clearance of the parasites on day 8 after withdrawing the drug pressure as seen in the microscopy assay.

| Drugs | Conc. of compounds | Parasites |  |  |  |  |
| --- | --- | --- | --- | --- | --- | --- |
|  |  | *B. bovis* | *B. bigemina* | *B. divergens* | *B. caballi* | *T. equi* |
| TC | 0.25×IC<sub>50</sub> | + | + | + | + | + |
|  | 0.5×IC<sub>50</sub> | + | + | + | + | + |
|  | 1×IC<sub>50</sub> | + | + | + | + | + |
|  | 2 ×IC<sub>50</sub> | - | - | + | - | + |
|  | 4 ×IC<sub>50</sub> | - | - | - | - | + |
| CH | 0.25×IC<sub>50</sub> | + | + | + | + | + |
|  | 0.5×IC<sub>50</sub> | + | + | + | + | + |
|  | 1×IC<sub>50</sub> | + | + | + | + | + |
|  | 2 ×IC<sub>50</sub> | + | + | + | + | + |
|  | 4 ×IC<sub>50</sub> | - | - | - | - | + |
|  | Untreated control | + | + | + | + | + |

Micrographs of TC- and CH-treated *B. bovis* **(Fig 4)**, *B. bigemina*, *B. divergens*, *B. caballi* **(Fig 5)**, and *T. equi* consistently showed degeneration of the parasites by loss of the typical shapes at 24 h, whereas further observations at 72 h showed deeply stained dot-shaped remnants of the parasites lodged within the erythrocytes.

**Fig 4.**
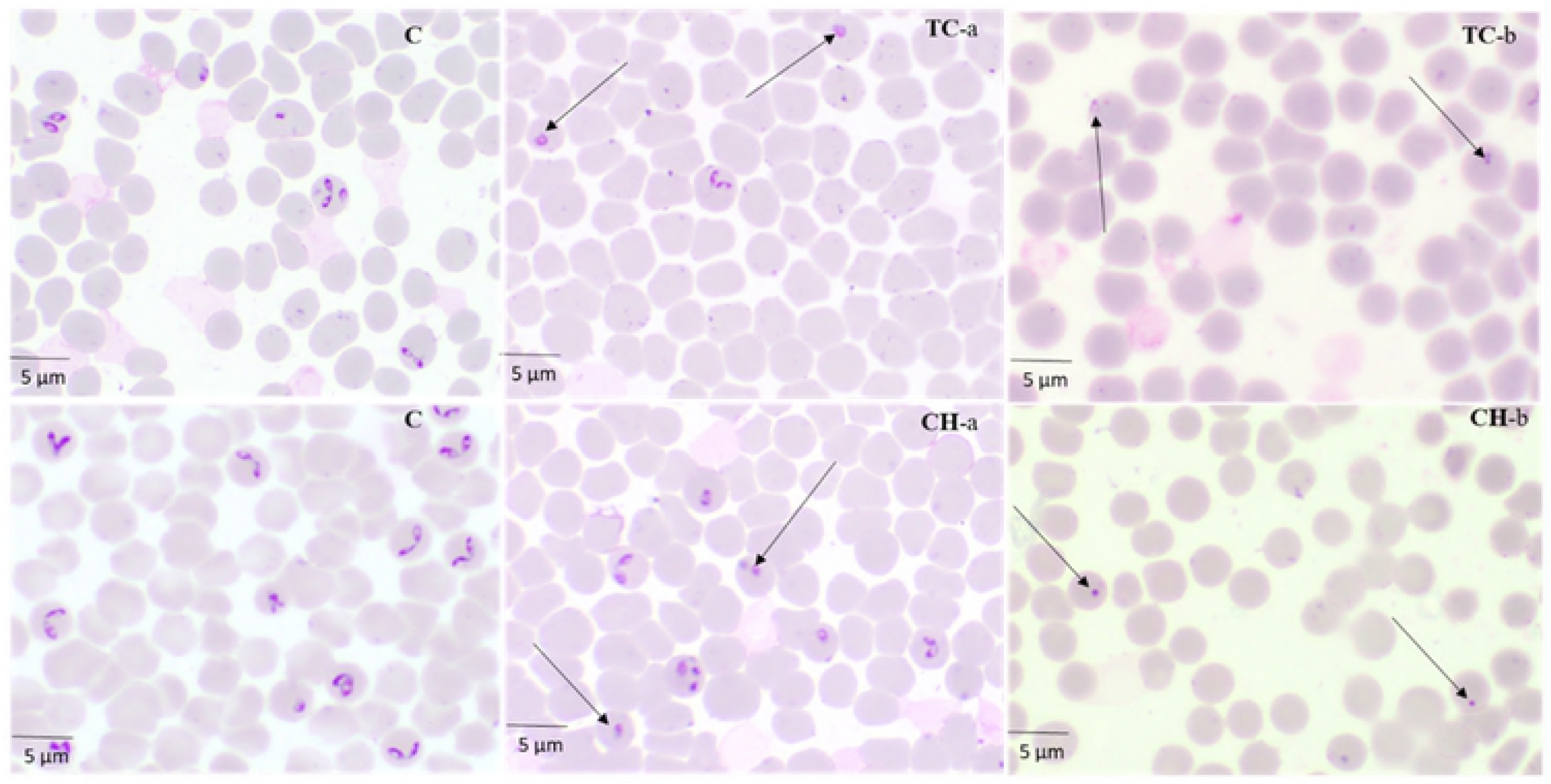
The morphological changes observed in TC- and CH-treated *B. bovis*. The arrows show TC- and CH-treated *B. bovis* parasites. The micrographs, C, were taken from the untreated wells. TC-a and CH-a were taken from treated wells at 24 h, while TC-b and CH-b were taken at 72 h.

**Fig 5.**
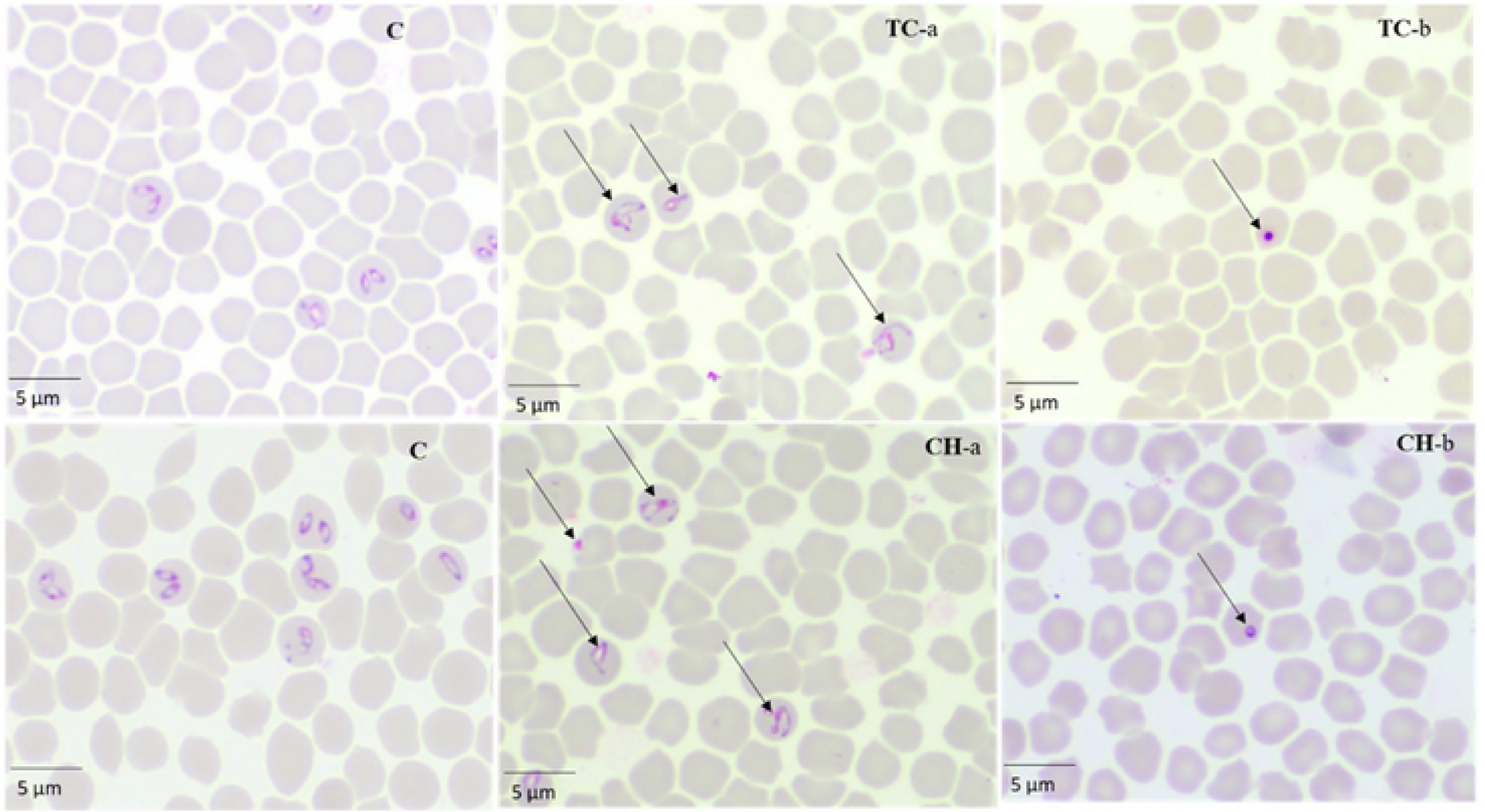
The morphological changes observed in TC- and CH-treated *B. caballi*. The arrows show TC- and CH-treated *B. caballi* parasites. The micrographs, C, were taken from the untreated wells. TC-a and CH-a were taken from treated wells at 24 h, while TC-b and CH-b were taken at 72 h.

### 3.3. The effects of a combination of trans-chalcone or chalcone hydrate with diminazene aceturate, atovaquone, or clofazimine *in vitro*

A drug-combination analysis was performed to determine whether the combined treatments are synergistic (have a greater effect), additive (have a similar effect), or antagonistic (have a reduced effect or block the effect). Five dilutions of CH or TC (**S2 Table**), as recommended in the Chou–Talalay method [25], were combined at a constant ratio with DA, AQ, or CF. The inhibition percentage for the single drug and each combination was analyzed using CompuSyn software to generate the combination index (CI) values (**S3Table**). The combination treatments of TC–DA and CH–DA showed an additive effect against *B. bovis* and *B. bigemina* and a synergistic effect against *B. divergens*, *B. caballi*, and *T. equi*. The combination treatments of TC–AQ showed a synergistic effect against *B. bigemina*, *B. caballi*, and *T. equi* and an additive effect against *B. bovis* and *B. divergens*. The combination treatments of CH–AQ showed a synergistic effect against *B. bovis*, *B. divergens*, *B. caballi*, and *T. equi* and an additive effect against *B. bigemina*. The combination treatments of TC–CF showed a synergistic effect against *B. bigemina*, *B. divergens*, *B. caballi*, and *T. equi* and an additive effect against *B. bovis.* The combination treatments of CH–CF showed a synergistic effect against *B. bigemina*, *B. caballi*, and *T. equi* and an additive effect against *B. bovis* and *B. divergens.* None of the combinations showed an antagonistic effect **(Table 3).**

**Table 3.** The effect of TC or CH with DA, AQ, or CF against *Babesia* and *Theileria parasites in vitro*

| Drug combinations |  | Parasites |  |  |  |  |
| --- | --- | --- | --- | --- | --- | --- |
|  |  | *B. bovis* | *B. bigemina* | *B. divergens* | *B. caballi* | *T. equi* |
| TC+DA | CI values | 1.05573 | 1.0588 | 0.51622 | 0.14276 | 0.11061 |
|  | Degree of association | Additive | Additive | Synergistic | Synergistic | Synergistic |
| CH+DA | CI values | 1.06140 | 1.0733 | 0.51622 | 0.20318 | 0.04491 |
|  | Degree of association | Additive | Additive | Synergistic | Synergistic | Synergistic |
| TC+AQ | CI values | 1.0912 | 0.07825 | 0.97607 | 0.22977 | 0.52253 |
|  | Degree of association | Additive | Synergistic | Additive | Synergistic | Synergistic |
| CH+AQ | CI values | 0.38743 | 1.09269 | 0.76130 | 0.07987 | 0.10921 |
|  | Degree of association | Synergistic | Additive | Synergistic | Synergistic | Synergistic |
| TC+CF | CI values | 1.03190 | 0.11538 | 0.72582 | 0.73026 | 0.84954 |
|  | Degree of association | Additive | Synergistic | Synergistic | Synergistic | Synergistic |
| CH+CF | CI values | 1.04538 | 0.16805 | 1.00823 | 0.32754 | 0.48165 |
|  | Degree of association | Additive | Synergistic | Additive | Synergistic | Synergistic |

CI denotes the weighted average combination index value.

### 3.4. Toxicity of *trans*-chalcone, chalcone hydrate, diminazene aceturate, atovaquone, and clofazimine on MDBK, NIH/3T3, and HFF cell lines

*Trans*-chalcone and chalcone hydrate showed an inhibitory effect on the *in vitro* culture of *Babesia* and *Theileria* parasites. Therefore, the effect of TC and CH on the host cells was evaluated using MDBK, NIH/3T3, and HFF cell lines to see the cytotoxicity of the two compounds **(Table 1)**. The EC_50_ values of TC on MDBK, NIH/3T3, and HFF cell lines were 293.9 ± 2.9, 434.4 ± 2.7, and 498 ± 3.1 µM, respectively. The EC_50_ values of CH on MDBK, NIH/3T3, and HFF cell lines were 252.7 ± 1.7, 406.3 ± 9.7, and 466 ± 5.7 µM, respectively **(Table 1)**. In a separate assay, DA and AQ at 100 µM did not show any inhibition of MDBK, NIH/3T3, or HFF cell viability, while CF showed inhibition only of MDBK with an EC_50_ value of 34 ± 3.4 µM **(S1 Table)**. The selectivity index, defined as the ratio of EC_50_ of the drugs tested on the cell line to IC_50_ of the tested drugs on *in vitro* culture of parasites. For TC, the highest selectivity index was achieved on *T. equi*, for the MDBK cell line the selectivity index was found to be 20.6 times higher than its IC_50_ on *T. equi*, while in the case of the NIH/3T3 cell line was found to be 30.4 times higher than the IC_50_ and in the case of the HFF cell line showed selectivity index 34.8 times higher than its IC_50_ on *T. equi*. For CH, the highest selectivity index was achieved on *T. equi* as in the case of the MDBK, NIH/3T3 and HFF cell lines was found to be 13.2 times, 21.2 times and 24.3 times higher than IC_50_, respectively **(Table 1).**

### 3.5. The chemotherapeutic effect of *trans*-chalcone against *B. microti* in mice

For further evaluation of TC efficacy in comparison with other drugs, the chemotherapeutic effect of TC was examined in mice infected with *B. microti* **(Fig 6)**.

**Fig 6.**
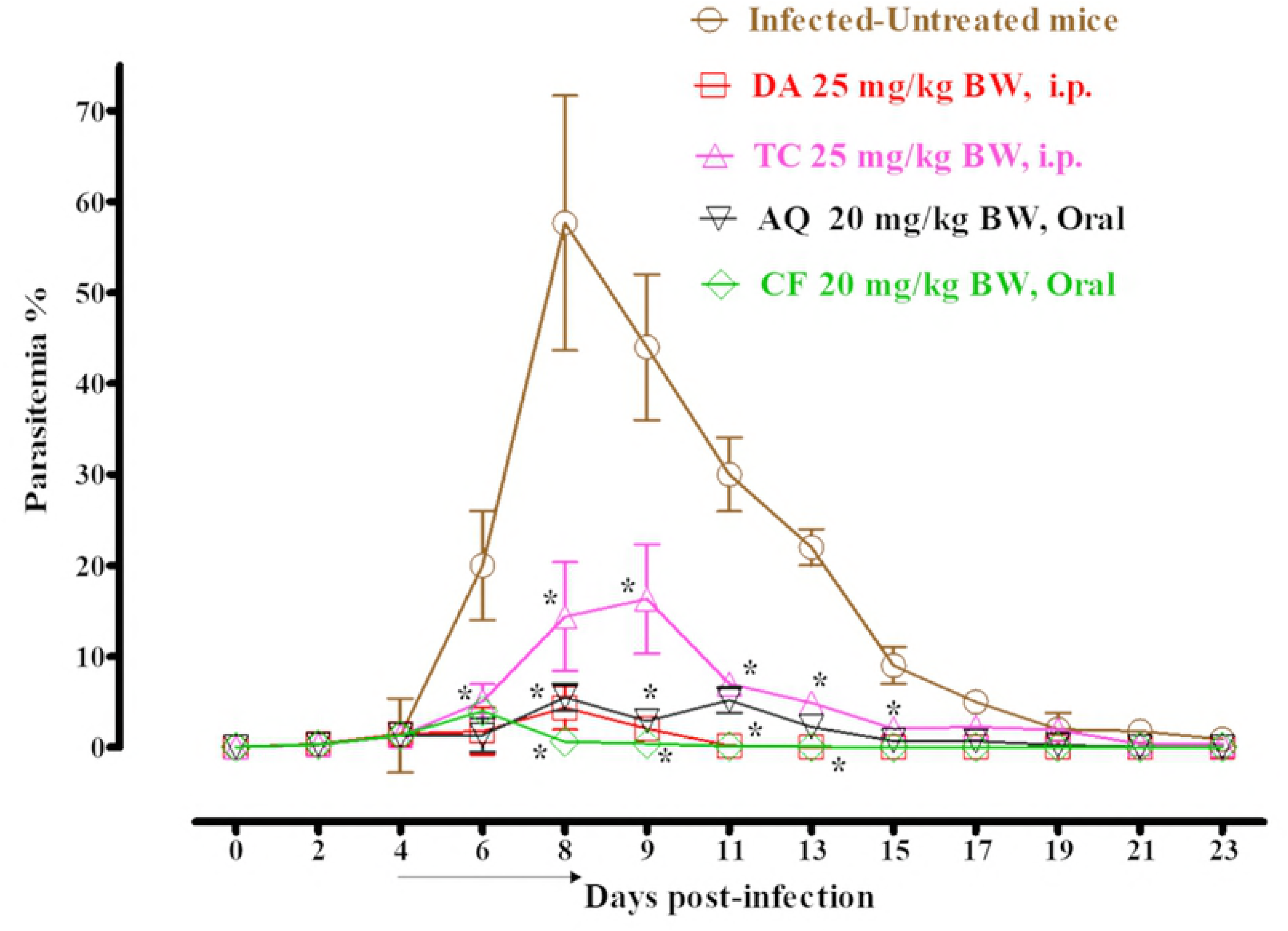
The growth inhibition of TC on *B. microti in vivo*. The graph shows the inhibitory effects of TC, DA, AQ, and CF treatments as compared with the untreated group. The values plotted indicate the mean ± standard deviation for two separate experiments. The asterisks (*) indicate statistical significance (*p* < 0.05) based on the unpaired *t*-test analysis. The arrow indicates 5 consecutive days of treatment. Parasitemia was calculated by counting infected RBCs among 2,000 RBCs using Giemsa-stained thin blood smears.

In the DDW control group, the multiplication of *B. microti* increased significantly and reached the highest parasitemia at 57.7% on day 8 post infection (p.i). In all treated groups, the level of parasitemia was cleared at a significantly lower percent of parasitemia than the control group (*p*< *0.05*) from days 6–12 p.i. In the monochemotherapy-treated mice, the peak parasitemia level reached 16.3% on day 9, 4.4% on day 8, 5.5% on day 8, and 4% on day 6 with 25 mg/kg TC, 25 mg/kg DA, 20 mg/kg AQ, and 20 mg/kg CF, respectively **(Fig 6)**. The parasitemia was undetectable via microscopy starting on day 13, 15, and 13 p.i. in mice treated with 25 mg/kg DA, 20 mg/kg AQ, and 20 mg/kg CF, respectively. In the combination-chemotherapy-treated groups, the peak parasitemia level reached 2.6%, 3.2%, and 10.4% with 12.5 mg/kg TC–12.5 mg/kg DA, 12.5 mg/kg TC–10 mg/kg CF, and 12.5 mg/kg TC–10 mg/kg AQ, respectively, on day 9 **(Fig 7)**.

**Fig 7.**
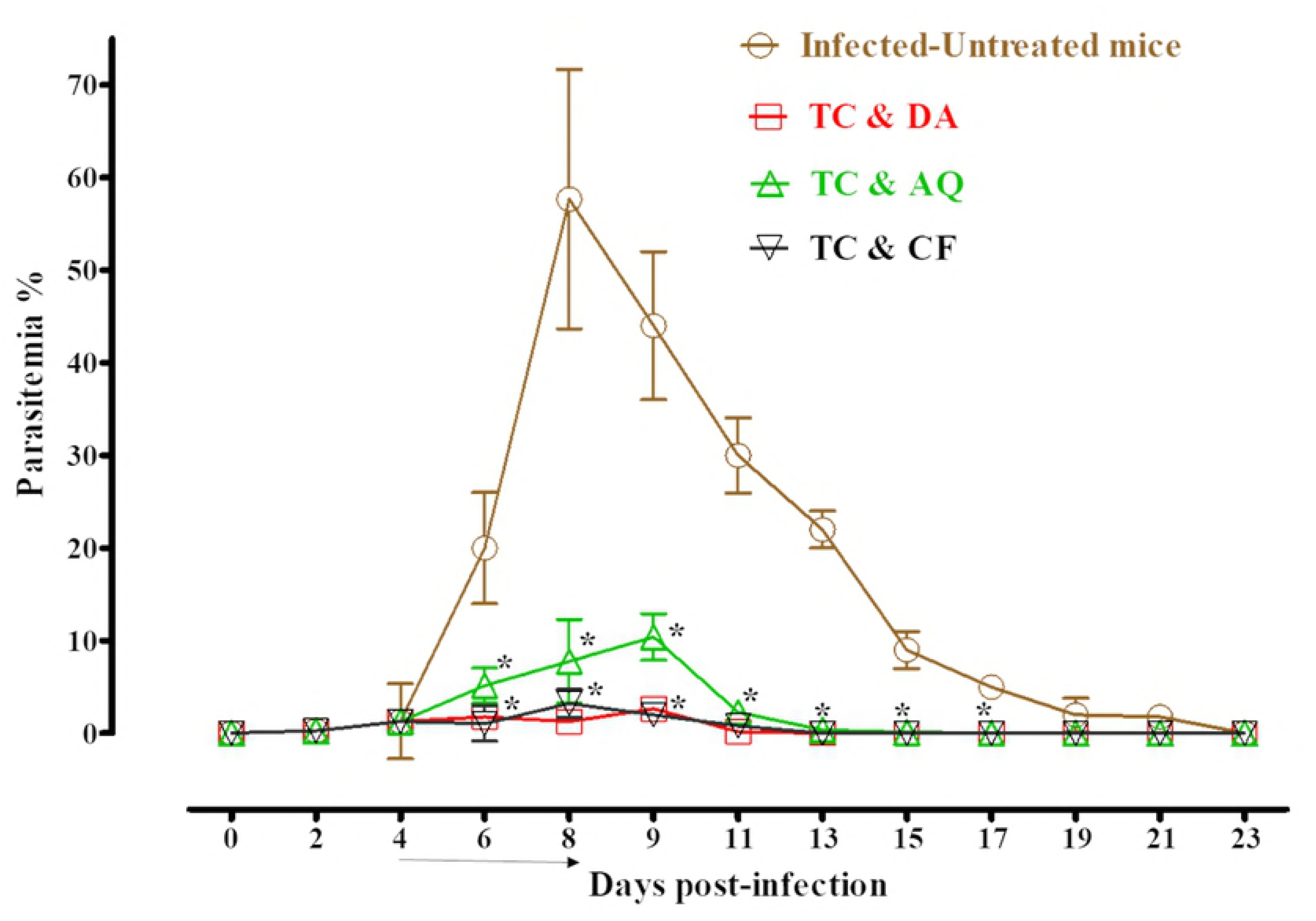
The growth inhibition of TC combinations on *B. microti in vivo*. The graph shows the inhibitory effects of DA, AQ, and CF combined with TC treatments as compared with the untreated group. The values plotted indicate the mean ± standard deviation for two separate experiments. The asterisks (*) indicate statistical significance (*p* < 0.05) based on the unpaired *t*-test analysis. The arrow indicates 5 consecutive days of treatment. Parasitemia was calculated by counting infected RBCs among 2,000 RBCs using Giemsa-stained thin blood smears.

The parasitemia was undetectable in microscopy examination on day 21 p.i. in mice treated with 25 mg/kg TC. The parasitemia was undetectable in mice via microscopy assay on days 13, 17, and 21 p.i. with 12.5 mg/kg TC–12.5 mg/kg DA, 12.5 mg/kg TC–10 mg/kg CF, and 12.5 mg/kg TC–10 mg/kg AQ, respectively. The parasite DNA was not detected on day 45 with 25 mg/kg DA IP, 12.5 mg/kg TC–10 mg/kg CF, or 12.5 mg/kg TC–12.5 mg/kg DA. In all other groups (20 mg/kg AQ oral, 20 mg/kg CF oral, 25 mg/kg TC IP, and 12.5 mg/kg TC–10 mg/kg AQ), the parasite DNA was detected until day 45 **(Fig 8)**.

**Fig 8.**
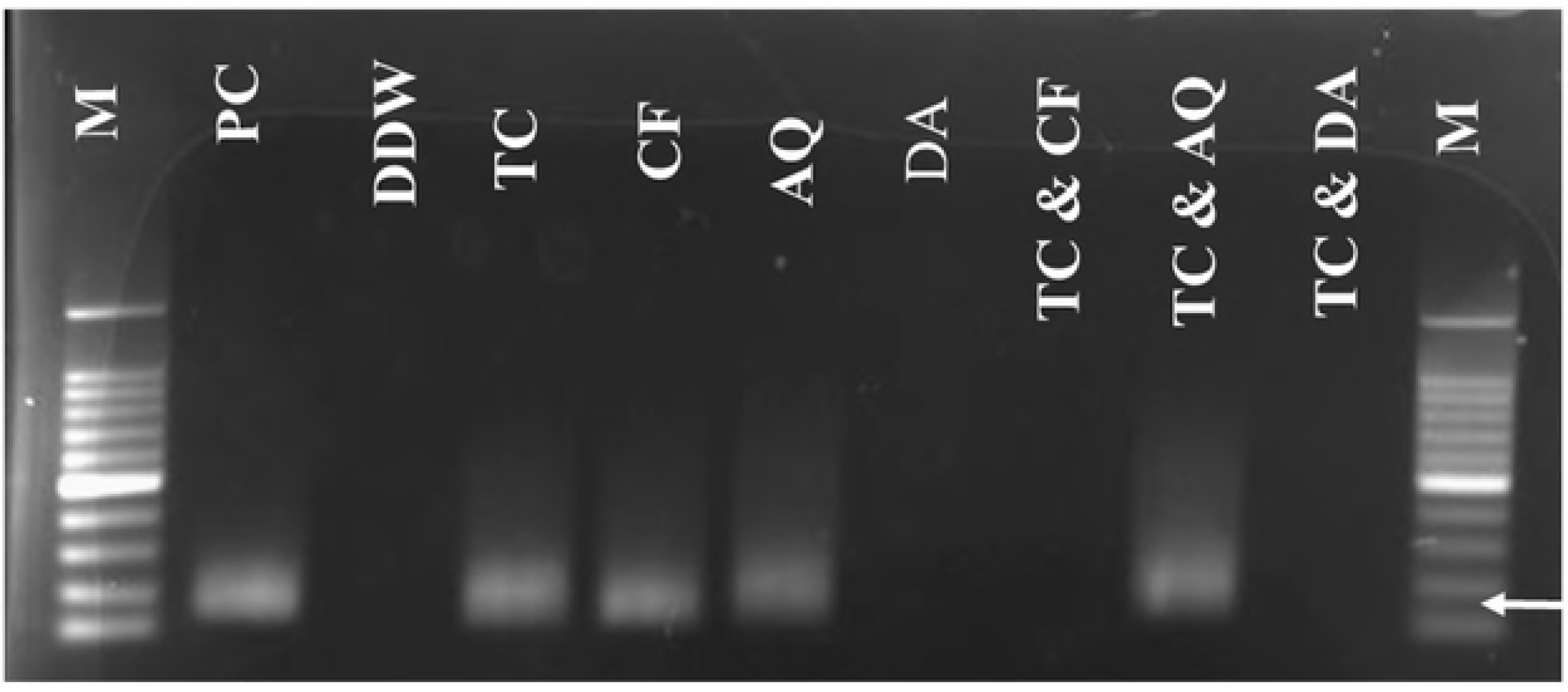
Molecular detection of parasite DNA in the treated groups. The image shows the molecular detection of parasites in the treated groups. The double distilled water (DDW) was used as a negative control, and M is for the marker. The arrow shows the expected band length of 154 bp for positive cases of *B. microti*.

Furthermore, infection with *B. microti* reduces the RBC count **(Fig 9A)**, hemoglobin concentration **(Fig 9B)**, and hematocrit percentage **(Fig 9C)** in mouse blood, as observed in the DDW control group on days 8 and 12 p.i. Significant differences (*p* < 0. 05) in RBC count were observed between the DDW control group and all drug-treated groups on days 8 and 12.

**Fig 9.**
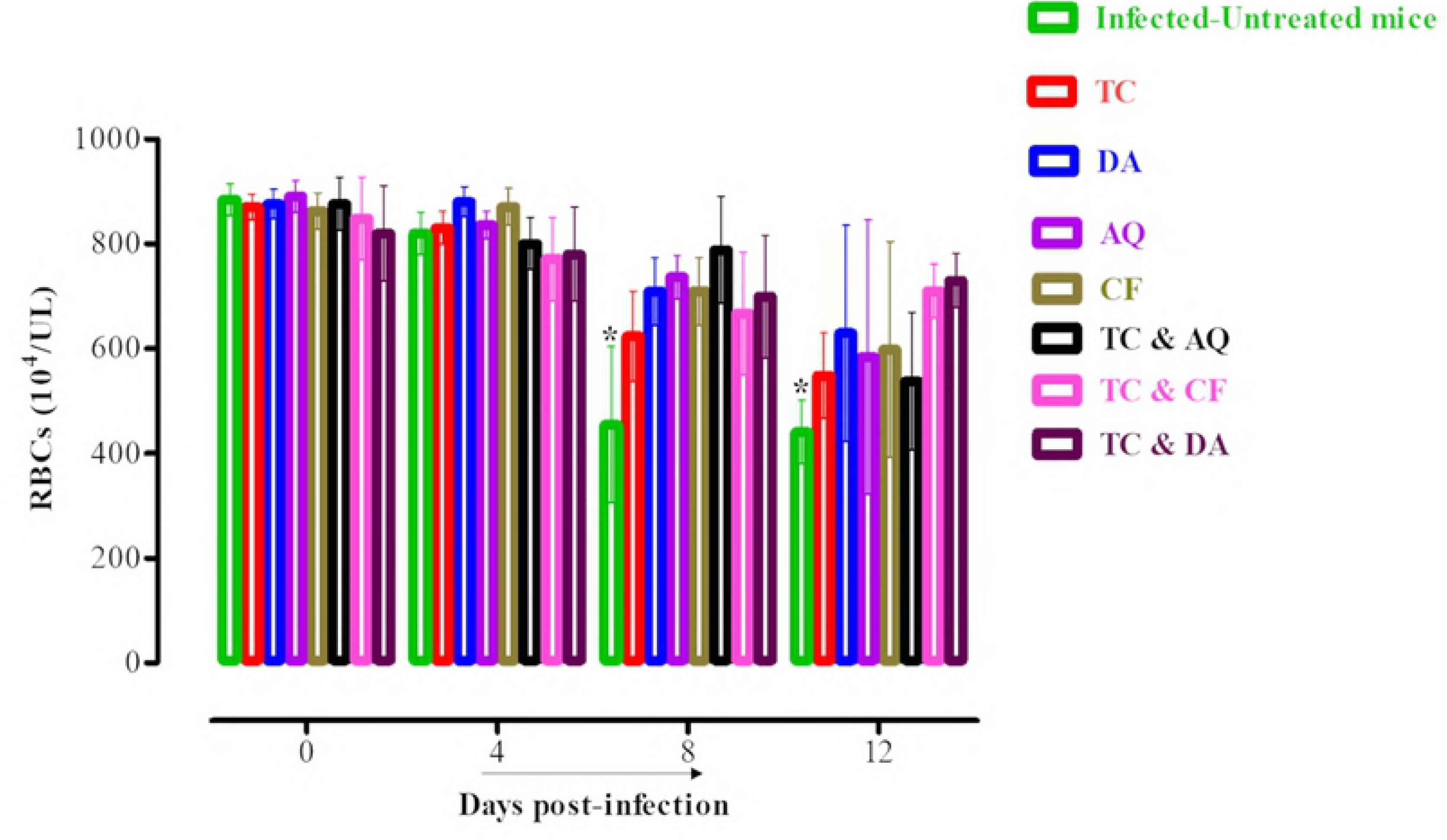
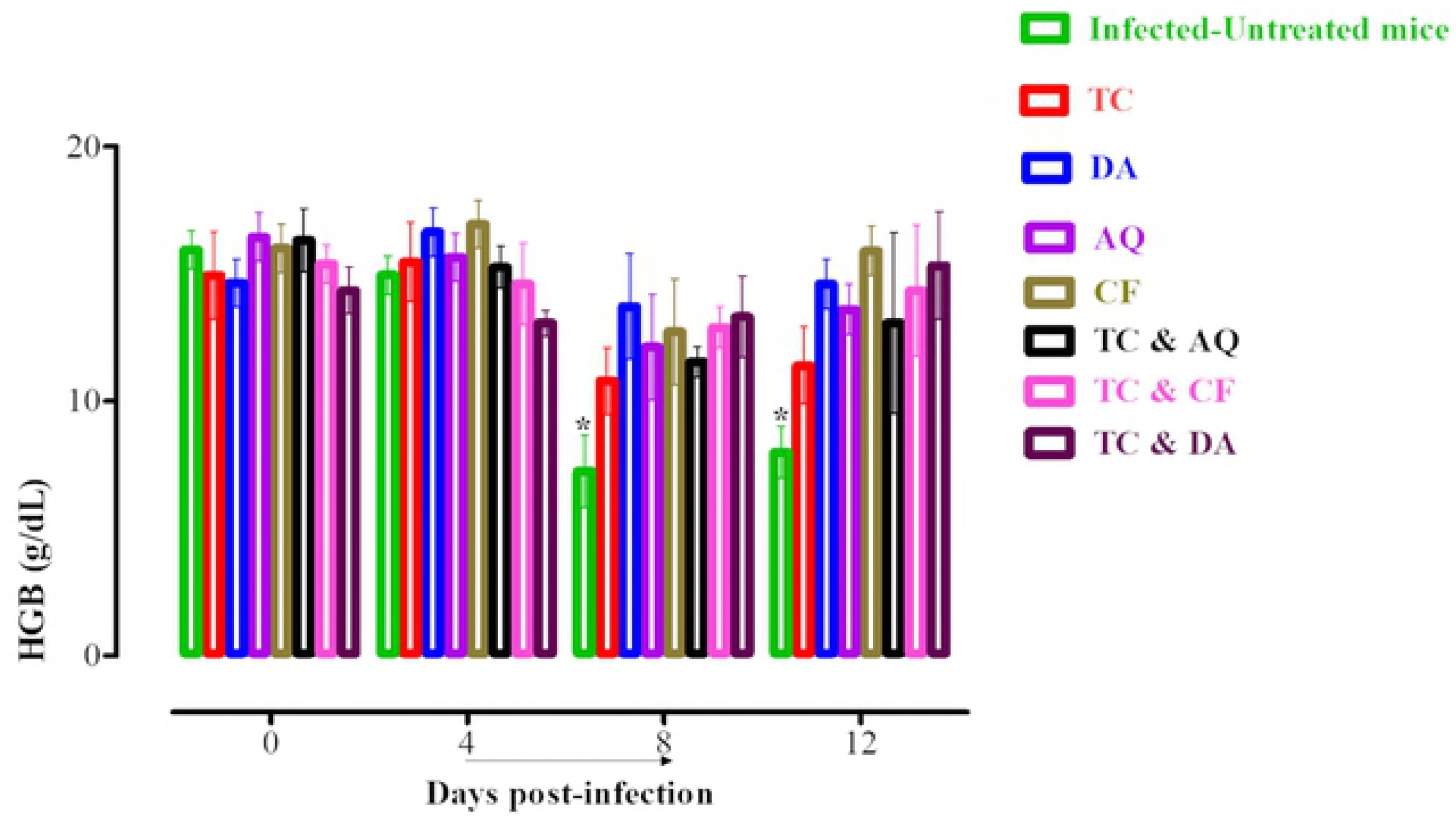
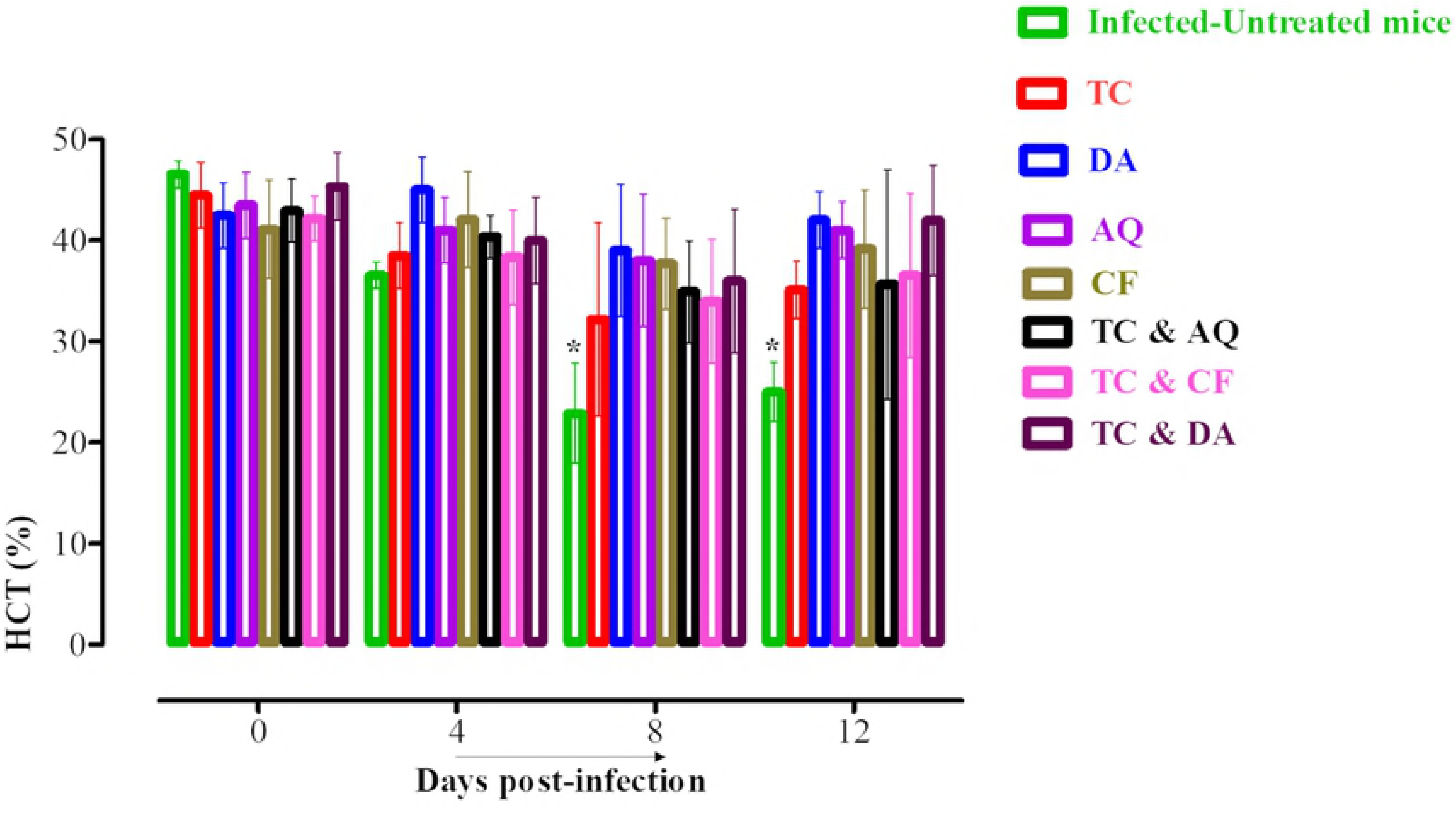
The changes in blood parameters in treated and untreated mice *in vivo*. The graphs show changes in the number of red blood cells (RBCs) (A), hemoglobin concentration (HGB) (B), and hematocrit percentage (HCT) (C) in different groups of treated mice as compared with untreated mice. The values plotted are the mean ± standard deviation for two separate trials. Each group contained five mice. The asterisks (*) indicate statistical significance (*p* < 0.05) based on the unpaired *t*-test analysis.

## 4. Discussion

The treatment of bovine and equine piroplasmosis is limited to diminazene aceturate (DA) and imidocarb propionate, while clindamycin–quinine and atovaquone–azithromycin combinations have been utilized to manage human babesiosis [8, 26]. Unfortunately, toxic effects and resistance of the piroplasms against the current drug molecules have been documented [2]. To overcome this challenge, research is urgently needed to discover new drug candidates and drug targets against piroplasms [8]. Therefore, the current study assessed the chemotherapeutic potential of CH and *trans*-chalcone (TC) against *Babesia* and *Theileria* parasites *in vitro* and *B. microti in vivo*. Further, the effects of combining CH and TC with the currently available drugs, namely, AQ, CF, and DA, against *Babesia* and *Theileria* parasites were assessed *in vitro*.

In the current study, both CH and TC were effective against the *Babesia* and *Theileria* parasites *in vitro* **(Figs 2 and 3)**. It is noteworthy that CH and TC were most effective against *T. equi*, followed by *B. caballi*, *B. bigemina*, and *B. divergens*, whereas they were least effective against *B. bovis*. The IC_50_ values shown by CH and TC against *Babesia* and *Theileria* parasites were comparable to those shown by CH and TC against *P. falciparum*, *Trypanosoma*, and *Leishmania* [9, 10, 13, 14, 22, 27]. This emphasizes that CH and TC are effective against many protozoan parasites. However, the mode of action has yet to be understood comprehensively in comparison with the existing data. In a previous study, Mi-Ichi et al. (2005) documented that chalcones are mitochondrial electron transport inhibitors that block ubiquinone (UQ) from binding to cytochrome *b* (*bc_1_*) in *Plasmodium* parasites and exhibit potent antimalarial activity[21]. Torres-Santos et al. (2009) and Chen et al. (2001) reported that chalcones inhibit the growth of *Leishmania* and *Trypanosomes* by inhibiting the activity of fumarate reductase (FRD), one of the enzymes of the parasite respiratory chain that it is very important in the energy metabolism of the parasites [12, 15]. Since this enzyme is absent from mammalian cells, it could be an important target for drugs against protozoan parasites. Based on the previous findings, it is possible that chalcones also inhibit the mitochondrial respiratory chain enzymes in *Babesia* and *Theileria* parasites, which could be elucidated in future studies.

The viability assay showed that TC and CH were more effective against *Babesia* parasites than against *Theileria* parasites. *B. bovis*, *B. bigemina*, and *B. caballi* could not relapse at 2×IC_50_ treatments of TC, while *B. divergens* could not relapse at 4×IC_50_ treatments of TC. *B. bovis*, *B. bigemina*, *B. caballi*, and *B. divergens* could not relapse at 4×IC_50_ treatments of CH. In contrast, *T. equi* recovered even at 4×IC_50_ treatments of CH and TC. This finding was similar to deductions by Tayebwa et al. (2018), who suggested that *T. equi* has better coping mechanisms than *Babesia* species [24]. However, the mechanism preventing *T. equi* from being completely killed by CH and TC remains unknown. In an attempt to visualize the morphological changes of CH- and TC-treated *Babesia* and *Theileria* parasites, micrographs were taken at various incubation times. The observations showed deformed and dividing parasites and irregular parasite shapes at 24 h and pyknotic remnants within the iRBCs at 72 h **(Figs 4 and 5)**. This showed that chalcones have a time-dependent effect on the *Babesia* and *Theileria* parasites. Although the exact mode of action is yet to be elucidated, the parasites progressively lost their shape and became smaller. This could be attributed to the ability of chalcones to interfere with the metabolic pathway, as documented in *P. falciparum* and *Leishmania* parasites [21, 28].

Combination chemotherapy has been recommended against drug-resistant protozoan and bacterial pathogens. Additionally, combination chemotherapy reduces drug dosages, thereby reducing their toxic side effects. Hence, the current study explored the combination of TC and CH with drugs such as DA, AQ, and CF *in vitro*. The findings of this study show that the effects of TC and CH combined with DA, AQ, or CF were either synergistic or additive against *Babesia* and *Theileria* parasites. The ability of TC and CH to combine with the current effective drugs is a property that can be explored in the development of chemotherapies against *Babesia* and *Theileria* [29]. That study showed that the CF–DA combination has additive effects on the *in vitro* growth of *B. bovis*, *B. bigemina*, and *B. caballi* and synergistic effects on that of *T. equi*, and the combination chemotherapy with low-dose regimens of CF and DA has a more potent inhibitory effect on *B. microti* in mice than did their monochemotherapies. It is imperative that further studies are performed to confirm the mechanisms of TC and CH against *Babesia* and *Theileria* so as to better understand the effect of interactions with other drugs such as DA, AQ, and CF.

The experiments to understand toxicity showed that CH and TC affected the viability of MDBK, NIH/3T3, and HFF cell lines with a dose-dependent inhibitory effect and a slightly high selectivity index. This finding is consistent with the results reported by Echeverria et al. (2009) [30]. They examined the cytotoxic activities of synthetic 2’-hydroxychalcones against hepatocellular carcinoma cells, demonstrating that synthetic 2’-hydroxychalcones show apoptosis induction and dose-dependent inhibition of cell proliferation without cytotoxic activities on normal cell lines [30]. Mi-Ichi et al. (2005) explained that the low cytotoxic activity of TC and CH against mammalian cell lines is attributed to the fact that the ubiquinol– cytochrome c reductase (UQCR) and succinate ubiquinone reductase (SQR) of *P. falciparum* mitochondria are different from those of the mammalian host cells [21]. With reference to the above, chalcones might be safe for use in animals and humans following further *in vivo* clinical studies.

The promising efficacy of TC *in vitro* prompted us to evaluate TC performance *in vivo*. TC administered intraperitoneally at a dose of 25 mg/kg resulted in a 71.8% inhibition of the parasitemia on day 9 p.i. However, the inhibition rate was lower than those in the presence of 25 mg/kg DA, 20 mg/kg AQ, and 20 mg/kg CF, which were 92.5%, 90.8%, and 93.1%, respectively **(Fig 6)**. Certainly, the additive and synergistic effects in these combinations were indicated by the high degree of association observed *in vitro*, which prompted studies *in vivo*. Therefore, the TC–DA and TC–CF combinations were evaluated in mice to determine whether combination treatment would enable the reduction of DA, AQ, and CF dosages without altering the therapeutic efficacy against *B. microti* infection. Interestingly, the combination treatment of TC and DA at a dose of 12.5 + 12.5 mg/kg improved the efficacy to 95.6%, while the combination of TC and AQ at a dose of 12.5 + 10 mg/kg resulted in 81.9% efficacy. The combination of TC and CF at a dose of 12.5 + 10 mg/kg resulted in a 94.4% inhibition in the parasitemia level at day 8 p.i. **(Fig 7)**. The potentiation of TC that was achieved in *in vivo* combination therapy confirms the result that was observed in the *in vitro* combination experiment, emphasizing that chalcones are good combinatorial drugs. With regard to the chemotherapeutic effects of TC against *Leishmania* in mice, Piñero et al. (2006) showed that a single dose of 4 mg/kg TC by subcutaneous administration could completely inhibit the pathogenicity of the *Leishmania* parasite *in vivo* [27]. In addition to being efficacious, chalcones enhanced the production of nitric oxide, which kills the intra-erythrocytic parasites and stimulates the host immune system [31].

In order to confirm the ability of TC to eliminate *B. microti*, a PCR assay was performed on samples collected on day 45 p.i. to be analyzed for the presence of DNA. Interestingly, this study confirmed the absence of *B. microti* DNA in groups treated with a combination chemotherapy of TC+DA or TC+CF **(Fig 8)** as compared to monotreatment. These results underscore the importance of combination chemotherapy in the effective control of piroplasmosis. This finding further emphasizes the need for combination therapy to achieve the most optimum efficacy and prevent the relapse of infection or development of a carrier state [29]. Furthermore, TC did not show toxic side effects to mice **(Fig 9A-C)**, consistent with a previous study [14]. Taken together, the findings advocate that TC is a potential drug against bovine and equine piroplasmosis.

## 5. Conclusion

CH and TC showed growth inhibitory and against *Babesia* and *Theileria in vitro.* Furthermore, TC showed chemotherapeutic efficacies against *B. microti in vivo*. TC effectiveness *in vivo* was comparable to that shown by DA, and it showed no toxicity to mice. The TC–DA and TC–CF combinations showed higher efficiency against piroplasms than did TC, DA, or CF monotherapies. This implies that TC could be used as a chemotherapeutic drug against piroplasmosis. Moreover, the results suggest that the TC–DA and TC–CF combination chemotherapies will be better choices for the treatment of piroplasmosis than TC, DA, or CF monotherapies.

## Acknowledgements

The authors would like to thank Dr. Bumduuren Tuvshintulga for assistance with sourcing chalcone hydrate and *trans*-chalcone.

## Supporting information

**S1 Table.** The IC_50_ and selectivity indices of AQ, DA, and CF (Control drugs).

**S1Table. The IC<sub>50</sub> and selectivity indices of AQ, DA, and CF (Control drugs).**
| Compound | Parasites | IC<sub>50</sub> ( μM )<sup>a</sup><br>parasites | EC<sub>50</sub> ( μM )<sup>b</sup> |  |  | Selective index <sup>c</sup> |  |  |
| --- | --- | --- | --- | --- | --- | --- | --- | --- |
|  |  |  | MDBK | NIH/3T3 | HFF | MDBK | NIH/3T3 | HFF |
| **AQ** | *B. bovis* | **0.039 ± 0.002** | **>100** | **>100** | **>100** | **> 10.4** | **> 10.4** | **> 10.4** |
|  | *B. bigemina* | **0.701 ± 0.04** |  |  |  | **> 12.7** | **> 12.7** | **> 12.7** |
|  | *B. divergens* | **0.038 ± 0.002** |  |  |  | **> 18.5** | **> 18.5** | **> 18.5** |
|  | *B. caballi* | **0.102 ± 0.0141** |  |  |  | **> 30.4** | **> 30.4** | **> 30.4** |
|  | *T. equi* | **0.095 ± 0.065** |  |  |  | **> 13.4** | **> 13.4** | **> 13.4** |
| **DA** | *B. bovis* | **0.35 ± 0.06** | **>100** | **>100** | **>100** | **> 285.7** | **> 285.7** | **> 285.7** |
|  | *B. bigemina* | **0.68 ± 0.09** |  |  |  | **> 208.3** | **> 208.3** | **> 208.3** |
|  | *B. divergens* | **0.43 ± 0.05** |  |  |  | **> 232.5** | **> 232.5** | **> 232.5** |
|  | *B. caballi* | **0.022 ± 0.0002** |  |  |  | **> 4545.5** | **> 4545.5** | **> 4545.5** |
|  | *T. equi* | **0.71 ± 0.05** |  |  |  | **> 476.2** | **> 476.2** | **> 476.2** |
| **CF** | *B. bovis* | **8.24 ± 1.7** | **34.7 ± 3.4** | **>100** | **>100** | **4.2** | **> 12.1** | **> 12.1** |
|  | *B. bigemina* | **5.73 ± 1.9** |  |  |  | **6.1** | **> 17.5** | **> 17.5** |
|  | *B. divergens* | **13.85 ± 4.3** |  |  |  | **2.5** | **> 7.2** | **> 7.2** |
|  | *B. caballi* | **7.95 ± 1.8** |  |  |  | **4.4** | **> 12.6** | **> 12.6** |
|  | *T. equi* | **2.88 ± 0.9** |  |  |  | **12.1** | **> 34.7** | **> 34.7** |

<sup>a</sup> Half maximum inhibition concentration of atovaquone (AQ), diminazene aceturate (DA), and clofazimine (CF) on the *in vitro* culture of parasites. The value was determined from the dose-response curve using non-linear regression (curve fit analysis). The values are the means of triplicate experiments.
<sup>b</sup> Half maximum effective concentration of AQ, DA, and CF on the cell line. The values were determined from the dose-response curve using non-linear regression (curve fit analysis). The values are the means of triplicate experiments.
<sup>c</sup> Ratio of the EC<sub>50</sub> of cell lines to the IC<sub>50</sub> of each species. High numbers are favorable.

**S2 Table.** Concentrations of *trans*-chalcone and chalcone hydrate combined with diminazene aceturate, atovaquone and clofazimine against *Babesia* and *Theileria* parasites *in vitro*.

| Parasite | Concentration (μM) | Trans-chalcone | Chalcone hydrate | Diminazine aceturate | Atovaquone | Clofazimine |
| --- | --- | --- | --- | --- | --- | --- |
| *B. bovis* | C<sub>1</sub> | 17.4 | 34.6 | 0.0875 | 0.00975 | 2.06 |
|  | C<sub>2</sub> | 34.8 | 69.2 | 0.175 | 0.0195 | 4.12 |
|  | C<sub>3</sub> | 69.6 | 138.4 | 0.35 | 0.039 | 8.24 |
|  | C<sub>4</sub> | 139.2 | 276.8 | 0.7 | 0.078 | 16.48 |
|  | C<sub>5</sub> | 278.4 | 553.6 | 1.4 | 0.156 | 32.96 |
| *B. bigemina* | C<sub>1</sub> | 8.33 | 15.225 | 0.17 | 0.17525 | 1.4325 |
|  | C<sub>2</sub> | 16.7 | 30.45 | 0.34 | 0.3505 | 2.865 |
|  | C<sub>3</sub> | 33.3 | 60.9 | 0.68 | 0.701 | 5.73 |
|  | C<sub>4</sub> | 66.6 | 121.8 | 1.36 | 1.402 | 11.46 |
|  | C<sub>5</sub> | 133.2 | 243.6 | 2.72 | 2.804 | 22.92 |
| *B. divergens* | C<sub>1</sub> | 16.2 | 20.575 | 0.1075 | 0.0095 | 3.4625 |
|  | C<sub>2</sub> | 32.4 | 41.15 | 0.215 | 0.019 | 6.925 |
|  | C<sub>3</sub> | 64.8 | 82.3 | 0.43 | 0.038 | 13.85 |
|  | C<sub>4</sub> | 129.6 | 164.6 | 0.86 | 0.076 | 27.7 |
|  | C<sub>5</sub> | 259.2 | 329.2 | 1.72 | 0.152 | 55.4 |
| *B. caballi* | C<sub>1</sub> | 4.725 | 6.975 | 0.0055 | 0.0255 | 1.9875 |
|  | C<sub>2</sub> | 9.45 | 13.95 | 0.011 | 0.051 | 3.975 |
|  | C<sub>3</sub> | 18.9 | 27.9 | 0.022 | 0.102 | 7.95 |
|  | C<sub>4</sub> | 37.8 | 55.8 | 0.044 | 0.204 | 15.9 |
|  | C<sub>5</sub> | 75.6 | 111.6 | 0.088 | 0.408 | 31.8 |
| *T. equi* | C<sub>1</sub> | 3.575 | 4.8 | 0.775 | 0.02375 | 0.72 |
|  | C<sub>2</sub> | 7.15 | 9.6 | 0.355 | 0.0475 | 1.44 |
|  | C<sub>3</sub> | 14.3 | 19.2 | 0.71 | 0.095 | 2.88 |
|  | C<sub>4</sub> | 28.6 | 38.4 | 1.42 | 0.19 | 5.76 |
|  | C<sub>5</sub> | 57.2 | 76.8 | 2.84 | 0.38 | 11.52 |

Note: <sup>a</sup> C<sub>1</sub>–C<sub>5</sub> refer to the concentrations (μM) 0.25×IC<sub>50</sub>, 0.5×IC<sub>50</sub>, 1×IC<sub>50</sub>, 2 ×IC<sub>50</sub>, 4 ×IC<sub>50</sub> of trans-chalcone, chalcone hydrate combined with diminazene aceturate, atovaquone and clofazimine. Combined concentrations were based on the calculated IC<sub>50</sub> values obtained from the *in vitro* fluorescence-based assay.

**S3 Table.** Calculation of weighted average of combination Index values.

| Parasites | Drug combinations <sup>a</sup> | Combination index values at |  |  |  | Weighted average<br>CI values <sup>b</sup> |
| --- | --- | --- | --- | --- | --- | --- |
|  |  | IC<sub>50</sub> | IC<sub>75</sub> | IC<sub>90</sub> | IC<sub>95</sub> |  |
| *B. bovis* | TC + DA | 0.7943 | 0.963 | 1.403 | 0.907 | 1.05573 |
|  | CH + DA | 2.710 | 0.625 | 0.882 | 1.002 | 1.06140 |
|  | TC + AQ | 0.614 | 0.991 | 0.768 | 1.503 | 1.0912 |
|  | CH + AQ | 0.4313 | 0.357 | 0.411 | 0.374 | 0.38743 |
|  | TC + CF | 0.2078 | 0.591 | 1.008 | 1.510 | 1.03190 |
|  | CH + CF | 0.3068 | 0.897 | 0.815 | 1.477 | 1.04538 |
| *B. bigemina* | TC +DA | 1.142 | 1.090 | 0.954 | 1.101 | 1.0588 |
|  | CH +DA | 1.580 | 1.047 | 1.005 | 1.011 | 1.0733 |
|  | TC +AQ | 0.4735 | 0.073 | 0.025 | 0.022 | 0.07825 |
|  | CH + AQ | 1.5459 | 0.777 | 1.209 | 1.050 | 1.09269 |
|  | TC +CF | 0.3658 | 0.011 | 0.118 | 0.103 | 0.11538 |
|  | CH + CF | 0.1765 | 0.293 | 0.010 | 0.222 | 0.16805 |
| *B. divergens* | TC +DA | 1.1862 | 0.359 | 0.418 | 0.501 | 0.51622 |
|  | CH +DA | 1.3292 | 0.572 | 0.319 | 0.433 | 0.51622 |
|  | TC +AQ | 1.3137 | 0.786 | 0.989 | 0.977 | 0.97607 |
|  | CH + AQ | 0.1360 | 0.981 | 0.701 | 0.853 | 0.76130 |
|  | TC +CF | 1.0332 | 0.541 | 0.777 | 0.703 | 0.72582 |
|  | CH + CF | 0.2613 | 0.810 | 1.303 | 1.073 | 1.00823 |
| *B. caballi* | TC +DA | 0.6646 | 0.118 | 0.165 | 0.008 | 0.14276 |
|  | CH +DA | 0.1238 | 0.052 | 0.372 | 0.172 | 0.20318 |
|  | TC +AQ | 0.9247 | 0.282 | 0.111 | 0.119 | 0.22977 |
|  | CH + AQ | 0.1077 | 0.103 | 0.091 | 0.053 | 0.07987 |
|  | TC +CF | 1.6806 | 0.697 | 0.608 | 0.601 | 0.73026 |
|  | CH + CF | 0.6604 | 0.192 | 0.113 | 0.473 | 0.32754 |
| *T. equi* | TC +DA | 0.0211 | 0.132 | 0.111 | 0.122 | 0.11061 |
|  | CH +DA | 0.3661 | 0.020 | 0.001 | 0.010 | 0.04491 |
|  | TC +AQ | 1.0503 | 0.537 | 0.431 | 0.452 | 0.52253 |
|  | CH + AQ | 0.0391 | 0.191 | 0.013 | 0.158 | 0.10921 |
|  | TC +CF | 1.5984 | 0.692 | 0.767 | 0.803 | 0.84954 |
|  | CH + CF | 0.9435 | 0.491 | 0.413 | 0.413 | 0.48165 |

CI value, combination index value; IC<sub>50</sub>, 50% inhibition concentration; DA, diminazene aceturate; AQ, atovaquone. <sup>a</sup> Two-drug combination between *trans*-chalcone, chalcone hydrate with diminazene aceturate, and atovaquone at a concentration of approximately 0.25 x IC<sub>50</sub>, 0.5 x IC<sub>50</sub>, IC<sub>50</sub>, 2 x IC<sub>50</sub>, and 4 x IC<sub>50</sub> (constant ratio). <sup>b</sup> The weighted average CI value was calculated with the formula [(1 x IC<sub>50</sub>) + (2 x IC<sub>75</sub>) + (3 x IC<sub>90</sub>) + (4 x IC<sub>95</sub>)]/10.

